# Reducing False Positives in CRISPR/Cas9 Screens from Copy Number Variations

**DOI:** 10.1101/247031

**Authors:** Alexander Wu, Tengfei Xiao, Teng Fei, X. Shirley Liu, Wei Li

## Abstract

CRISPR/Cas9 knockout screens have been widely used to interrogate gene functions across a wide range of cell systems. However, the screening outcome is biased in amplified genomic regions, due to the ability of the Cas9 nuclease to induce multiple double-stranded breaks and strong DNA damage responses at these regions. We developed algorithms to correct biases associated with copy number variations (CNV), even when the CNV profiles are unknown. We demonstrated that our methods effectively reduced false positives in amplified regions while preserving signals of true positives. In addition, we developed a sliding window approach to estimate regions of high copy numbers for cases in which CNV information is not available. These copy number estimations can subsequently be used to effectively correct CNV-related biases in CRISPR screening experiments. Our approach is integrated into the existing MAGeCK/MAGeCK-VISPR analysis pipelines and provides a convenient framework to improve the precision of CRISPR screening results.

## Introduction

The CRISPR/Cas9 (clustered regularly interspaced short palindromic repeats/CRISPR-associated9) system has been shown to be a highly effective genome editing tool for mammalian cells [1-3]. This system has led to the application of CRISPR/Cas9 loss-of-function genome screening [4, 5], in which tens of thousands of genes are knocked out and evaluated for association with cell proliferation (or other phenotypes). While CRISPR screens have demonstrated great promise for investigating gene functions in cancer and other research areas [6, 7, 8], the outcomes of CRISPR/Cas9 knockout screens are influenced by the copy number variations (CNVs) of regions targeted by single guide RNAs (sgRNAs) [9]. In regions with high CNVs, sgRNAs direct Cas9 to induce cuts at every single copy, triggering a stronger DNA damage response, cell cycle arrest, and decreasing cell proliferation. This leads to sgRNA depletion in screening readouts, even if the function of targeted regions is unrelated to the screening phenotype [10]. This problem is particularly relevant for cancer cells, as copy number alterations are common in human cancers [11]. Therefore, reducing or correcting these copy number-related effects is critical for improving the precision of downstream CRISPR screening analysis.

Currently, CERES is the only known computational method to decouple the copy number bias in CRISPR screens [12]. It models sgRNA readouts as the combination of both CNV bias and underlying gene knockout effects and uses a constrained least square optimization model to decouple both effects. While this approach is effective in reducing the effect of copy number bias in a large number of cell lines, it does not take into consideration many of the other biases that exist in CRISPR screen analyses, including frequent absence of replicates, variability in sgRNA knockout efficiencies, and variability in read count distributions. Furthermore, the model requires CNV profiles for each cell type as input and, thus, cannot be applied to cells with unknown CNV profiles.

We introduced two computational correction methods to correct copy number biases in CRISPR screens. For cells with known CNV profiles, we built a regression model to capture the relationship between CNV and gene essentiality, and subsequently correct the effects of CNV biases. We demonstrated that our method is effective in reducing the prevalence of false positives in high copy number regions, while retaining robust results for the identification of essential genes with CRISPR screens. In addition, we developed a computational model to approximate the relative CNVs for genomic regions that are susceptible to copy number biases. It operates by locally analyzing sgRNA read count changes over a broad genomic region, and subsequently estimating the copy number status based on the behaviors of all sgRNAs in the region. This approach is particularly suitable for screening experiments in which CNV profiles are unknown. We demonstrated that this approach correctly identified known amplified regions. Both methods are integrated into MAGeCK [13] and MAGeCK-VISPR [14], two computational algorithms we previously developed for the analysis of CRISPR screens.

## Methods

### MAGeCK and MAGeCK-VISPR

MAGeCK and MAGeCK-VISPR are algorithms we developed to estimate the effects of gene knockouts in CRISPR screens. MAGeCK builds a mean-variance model to estimate the variance of the read counts, and uses these variance estimations to model the read count changes for each sgRNA in the treatment samples relative to the control samples. The read count changes (calculated as “sgRNA scores” in MAGeCK) of all sgRNAs targeting each gene are then ranked and summarized into one score for the gene (“gene score”), using a modified robust ranking aggregation (RRA) algorithm. For clarification, we refer to this approach as “MAGeCK-RRA”.

In contrast, MAGeCK-VISPR consists of an algorithm named “MAGeCK-MLE” that estimates the essentiality of genes in a CRISPR screen via a maximum likelihood approach. MAGeCK-MLE initially takes in as input a raw table of reads, and the read count of each sgRNA for each sample is modeled as a negative binomial (NB) random variable. For each sgRNA *i* in sample *j*, the mean of the NB random variable is subsequently modeled as *μ_ij_ ∼* exp(*β_i0_ + ∑_r_ d_jr_ β_gr_*),where *β_i0_* represents the initial abundance of sgRNA *i*, and *β_gr_* describes the effect of knocking out targeting gene *g* for condition *r*. Importantly, *β_gr_* (named the “*β* score”) represents the essentiality of gene *g* in condition *r* and can be interpreted similarly to a log-fold change value: a negative *β* score suggests that gene *g* is negatively selected in condition *r*, and *vice versa*. The value of *β_gr_* is inferred by maximizing the joint log-likelihood of observing the read counts associated with all sgRNAs targeting gene *g* for condition *r*.

### CNV Bias Correction for CRISPR Screen Experiments using MAGeCK-MLE

To correct for copy number biases in MAGeCK-MLE, we model the relationship between *β* scores and copy numbers for all genes in condition *r* with the following equation:

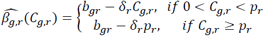

where *C_gr_* is the copy number estimation of gene *g*, *δ_r_* is a constant describing the relative effect of CNV profiles on *β* score, *p_r_* is the copy number threshold at which the cell cycle arrest is fully activated, and *b_gr_* is the *β* score for a gene whose knockout does not trigger any DNA damage response mechanism (*i.e.*, no CNV effect). The values of *b_gr_*, *p_r_*, and *δ_r_* are estimated for each condition *r* by minimizing the least squared error between the observed *β* scores and the scores *β* scores estimated by the equation defined with these parameters. An example of the model defining the relationship between *β* scores and gene copy numbers for MCF7 cells is presented in Figure 1a.

**Figure 1.**
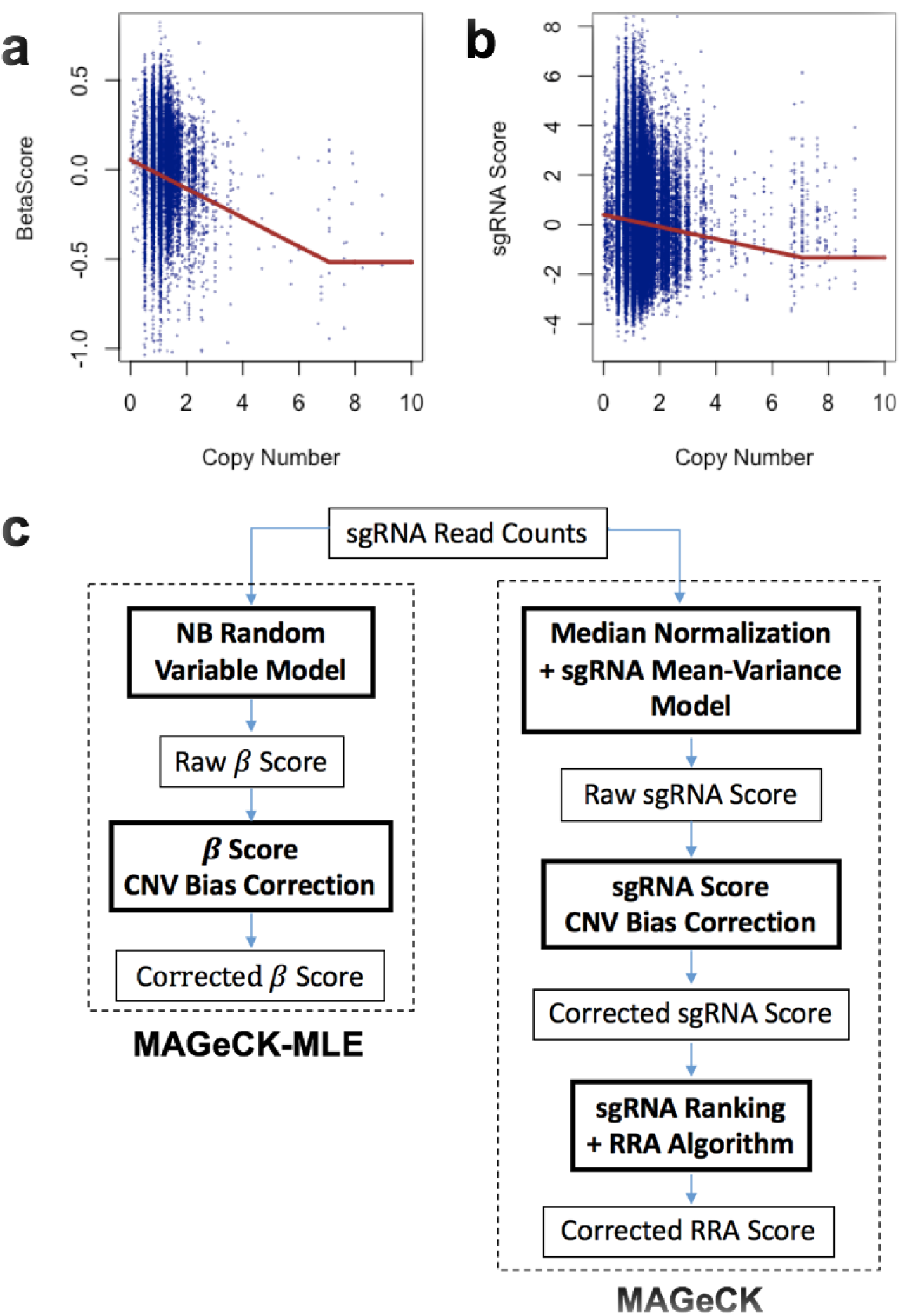
CNV Bias Modeling and Correction in MAGeCK-MLE and MAGeCK for MCF7. **(a-b)** Relationship between **(a)** *β* scores and gene copy numbers, and **(b)** sgRNA scores and gene copy numbers for the MCF7 cell line. Each blue point corresponds to the *β* (or sgRNA) score and copy number for a single gene. The function in red corresponds to the modeled relationship between all *β* (or sgRNA) scores and gene copy numbers for MCF7. **(c)** Workflow of the MAGeCK-MLE and MAGeCK algorithms with the inclusion of the CNV bias correction feature.

After defining the *β* score-copy number model, the *β* score for each gene *g* in condition *r* is adjusted by an additive correction factor Δ. The value of Δ is equivalent to the magnitude of the bias associated with the gene’s copy number relative to the bias associated with a gene of copy number 2. In other words, the adjusted *β* scores for gene *g* in condition *r* is defined as 
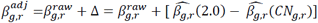
.

### CNV Bias Correction for CRISPR Screen Experiments using MAGeCK-RRA

In the MAGeCK-RRA algorithm, we correct for CNV biases in the sgRNA scores, or the sgRNA read count changes between two conditions (as calculated via the NB model). Similar to the MAGeCK-MLE CNV correction model, we model sgRNA scores and gene CNV profiles as follows:

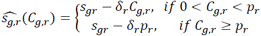

where *s_i,r_* is the score of sgRNA *i* in comparison *r*. An example of the sgRNA score and copy number variation is shown in Figure 1b. The sgRNA scores are similarly corrected for the copy number bias effect by introducing the additive corrective factor Δ to each sgRNA score: 
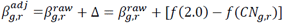
. After adjusting the sgRNA scores, the sgRNAs are ranked by their scores, and gene essentiality is estimated via the RRA algorithm. An example of the model defining the relationship between sgRNA scores and gene copy numbers for MCF7 cells is presented in Figure 1b.

A schematic of the CNV correction procedure for both MAGeCK-RRA and MAGeCK-MLE is presented in Figure 1c.

### CNV Estimation using sgRNA Read Counts from CRISPR Screen Experiments

The method mentioned above and other methods (like CERES) require CNV profiles as an input. In the absence of relevant copy number data, we estimate the relative gene copy numbers via a sliding window approach, in which we model sgRNA abundance changes within large genomic regions. This approach uses a 2 Mb window and a step size of 0.1 Mb as default settings to scan across entire chromosomes for all chromosomes in the genome. For each window, the log-fold change values associated with all sgRNAs in the window are aggregated. The mean of these values is then assigned as a *window score* if the corresponding window encompass 5 or more genes. Once all window scores are computed, each gene is assigned a *gene score w_g,r_*, that equals the mean of the window scores corresponding to the windows that overlap the gene. The relative copy number for each gene *g* in condition *r* is then calculated from the distribution of gene scores as follows: 
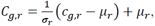
 where *c_g,r_* = −*w_g,r_* is the negated gene score, *C_g,r_* is the estimated relative copy number, *σ_r_* is the standard deviation of the distribution of the *c_g,r_* values, and *µ_r_* is the mean of the *c_g,r_* values. *σ_r_* is introduced to adjust for the variability in essentiality scores across different cell lines (i.e. some cell lines will have more negative/positive essentiality scores than others).

The relative copy number estimates can subsequently be incorporated into the copy number bias correction algorithms described above. In order to account for the strongly amplified genomic regions, only the sgRNA scores or *β* scores of the genes with the highest 2% of copy number estimates are adjusted.

## Results and Discussion

### CNV Bias Correction Reduces False Positives in Gene Essentiality Identification

We first applied the correction algorithms of MAGeCK-MLE and MAGeCK-RRA to CRISPR screens performed on MCF7 and T47D, two breast cancer cell lines. The MCF7 cell line possesses a high copy number region in Chromosome 17 that does not exist in the T47D cell line (Figure 2a). As expected, a high correlation was observed between amplified regions and strongly negative *β* scores (calculated by MAGeCK-MLE) and RRA scores (calculated by MAGeCK-RRA) before adjustment (Figure 2a). MAGeCK-MLE decreased the magnitude of *β* scores in these regions, reducing the effects of amplifications. Similar corrections to RRA scores were observed in MCF7 (Figure 2b). As for T47D, the raw and adjusted *β* (and RRA) scores showed little difference due to the absence of high copy numbers within this same region (Figures 2c, d).

**Figure 2.**
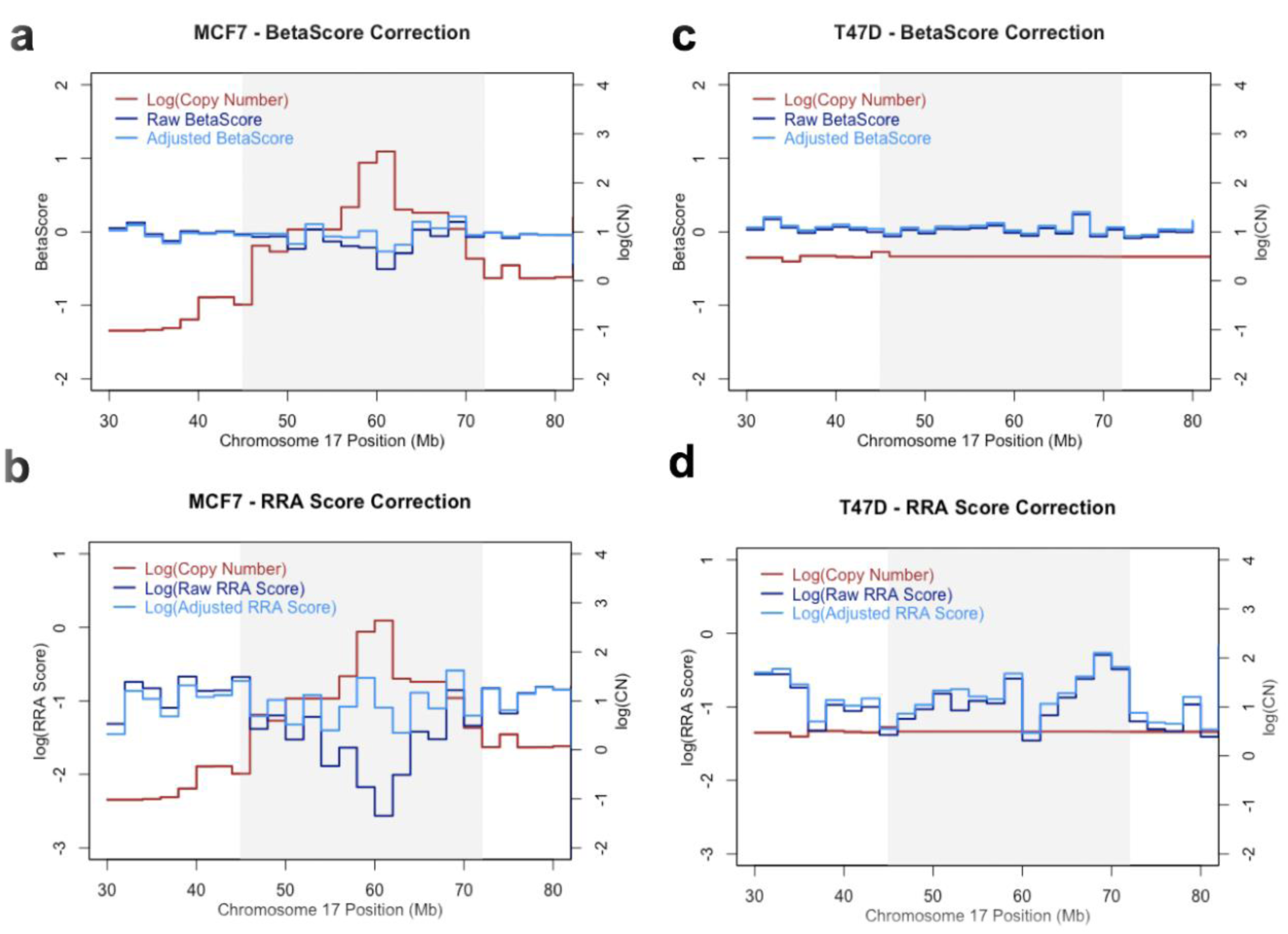
Essentiality Score Adjustments in a Variable Copy Number Region in Chromosome 17. High CN regions are highlighted in gray. **(a)** *β* scores in a high CN region in MCF7. **(b)** log(RRA scores) in a high CN region in MCF7. **(c)** *β* scores in a low CN region in T47D. **(d)** log(RRA scores) in a low CN region T47D.

We systematically evaluated the relationship between copy numbers and CRISPR screening results in T47D and MCF7. Genes that have the highest copy numbers demonstrated significantly smaller *β* scores relative to the remaining genes in both cell lines (p < 2.2e-16 and p = 1.085e-9, respectively, using the Kolmogorov-Smirnov test). Upon inclusion of the CNV correction feature in MAGeCK-MLE, the difference in the distribution of *β* scores between the two groups of genes is smaller (p = 1.999e-4 and p = 0.5058 for MCF7 and T47D, respectively; Figure 3a-b). The same reduction of differences between high CNV genes and other genes is observed for RRA scores as well (Figure 3c-d).

**Figure 3.**
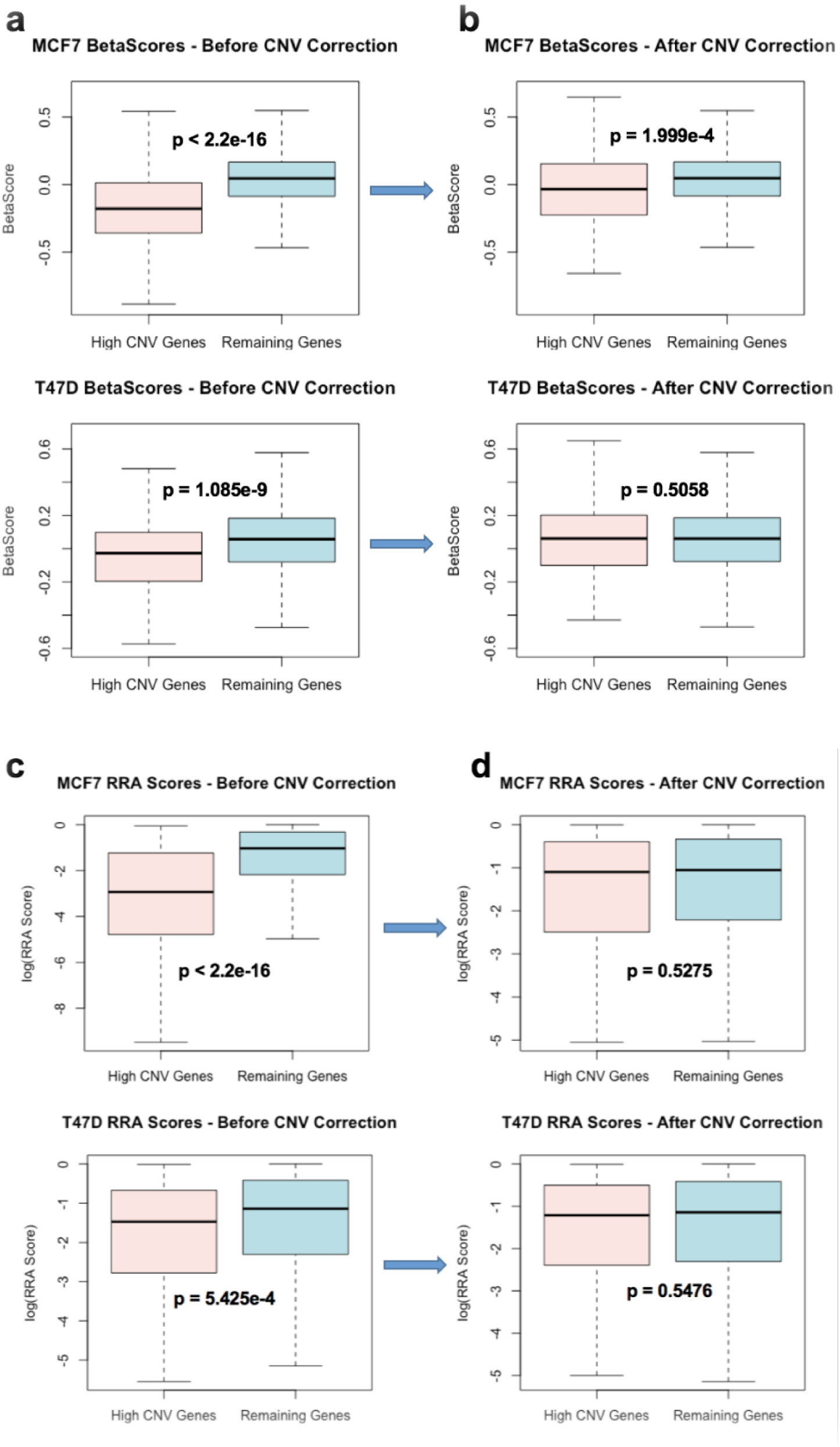
Comparison of Essentiality Score Distributions Between High Copy Number Genes and Remaining Genes. “High CNV genes” are the set of genes with the highest 1% of copy numbers. **(a-b)** Distribution of *β* scores from MAGeCK-MLE without copy number bias correction **(a)** and with copy number bias correction **(b)**. **(c-d)** Distribution of log(RRA scores) from MAGeCK without copy number bias correction **(c)** and with copy number bias correction **(d)**.

Gene set enrichment analysis (GSEA) also demonstrated that genes with high copy numbers are less enriched after CNV bias correction in negatively selected gene sets for MCF7 cells (Figure 4). Genes in amplified regions are less enriched after bias correction in T47D cells as well.

**Figure 4.**
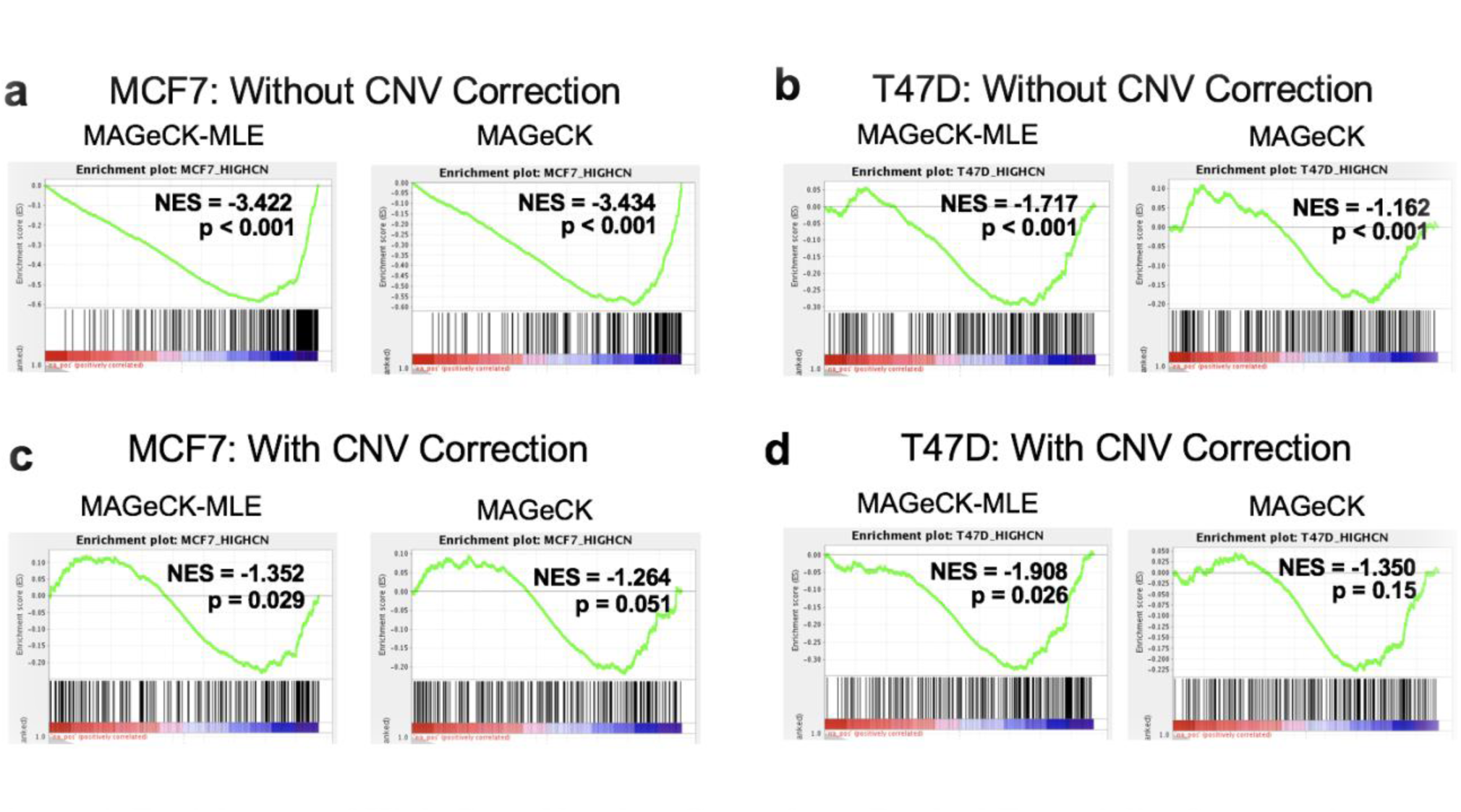
Enrichment of High Copy Number Genes from Ranked Essentiality Scores using Gene Set Enrichment Analysis (GSEA). **(a-b)** Enrichment of high CN genes in MCF7 based on the gene rankings before **(a)** and after **(b)** CNV correction. **(c-d)** Enrichment of high CN genes in T47D on the gene rankings before **(c)** and after **(d)** CNV correction. NES: Normalized Enrichment Score.

### CNV Bias Correction Retains Robust Results from MAGeCK-MLE and MAGeCK

We next tested whether the correction algorithms affected the identification of essential genes, by examining known essential genes in the amplified region of Chromosome 17. PPM1D, BRIP1, and PECAM1 are three known oncogenes in breast cancer cells that are frequently amplified in Chromosome 17 [15-19]. These genes are strongly negatively selected in the amplified region of MCF7 cells and remained strongly negatively selected after CNV correction, an indication that both correction algorithms do not eliminate truly essential genes in the screen (Figure 5a-b). In contrast, these genes only demonstrated moderate negative selection in T47D cells, which do not have amplifications in the corresponding genomic region (Figure 5c-d).

**Figure 5.**
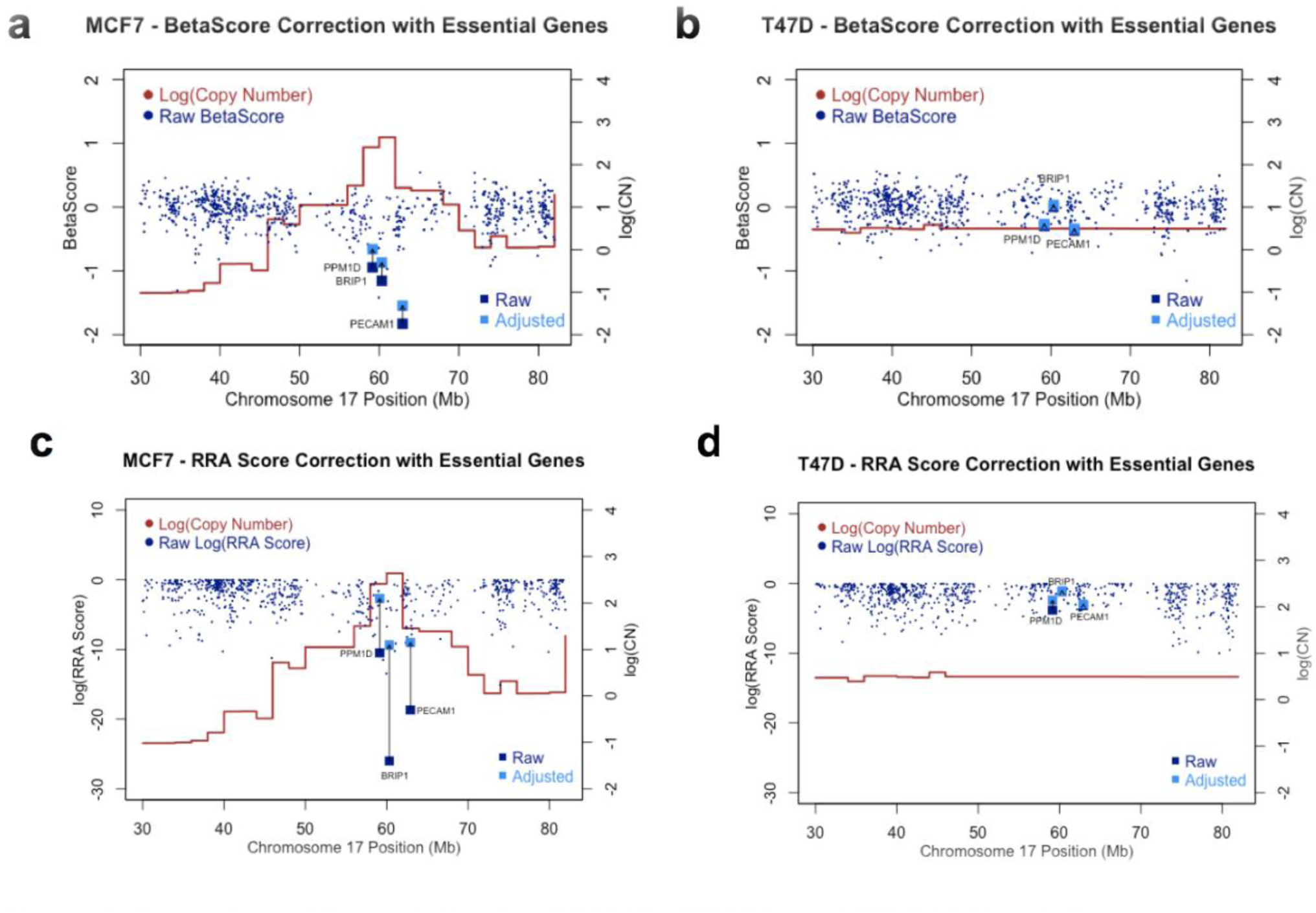
Retention of Essentiality for PPM1D, BRIP1, and PECAM1. **(a)** *β* scores adjustments in MCF7. **(b)** *β* score adjustments in T47D. **(c)** log(RRA score) adjustments in MCF7. **(d)** log(RRA score) adjustments in T47D.

To comprehensively evaluate the overall effects of CNV correction across the genome, we examined the enrichment of known essential genes in the negatively selected gene list. These genes include genes in the pathways associated with ribosomes and spliceosomes, both are known to be highly essential across the vast majority of cells. In both the MCF7 and T47D samples, ribosomes and splicesomes (either ranked by *β* scores or RRA scores) remain strongly enriched in essential gene list before and after correction (Figure 6), providing evidence that our algorithms preserve the signals from truly essential genes identified in CRISPR screens.

**Figure 6.**
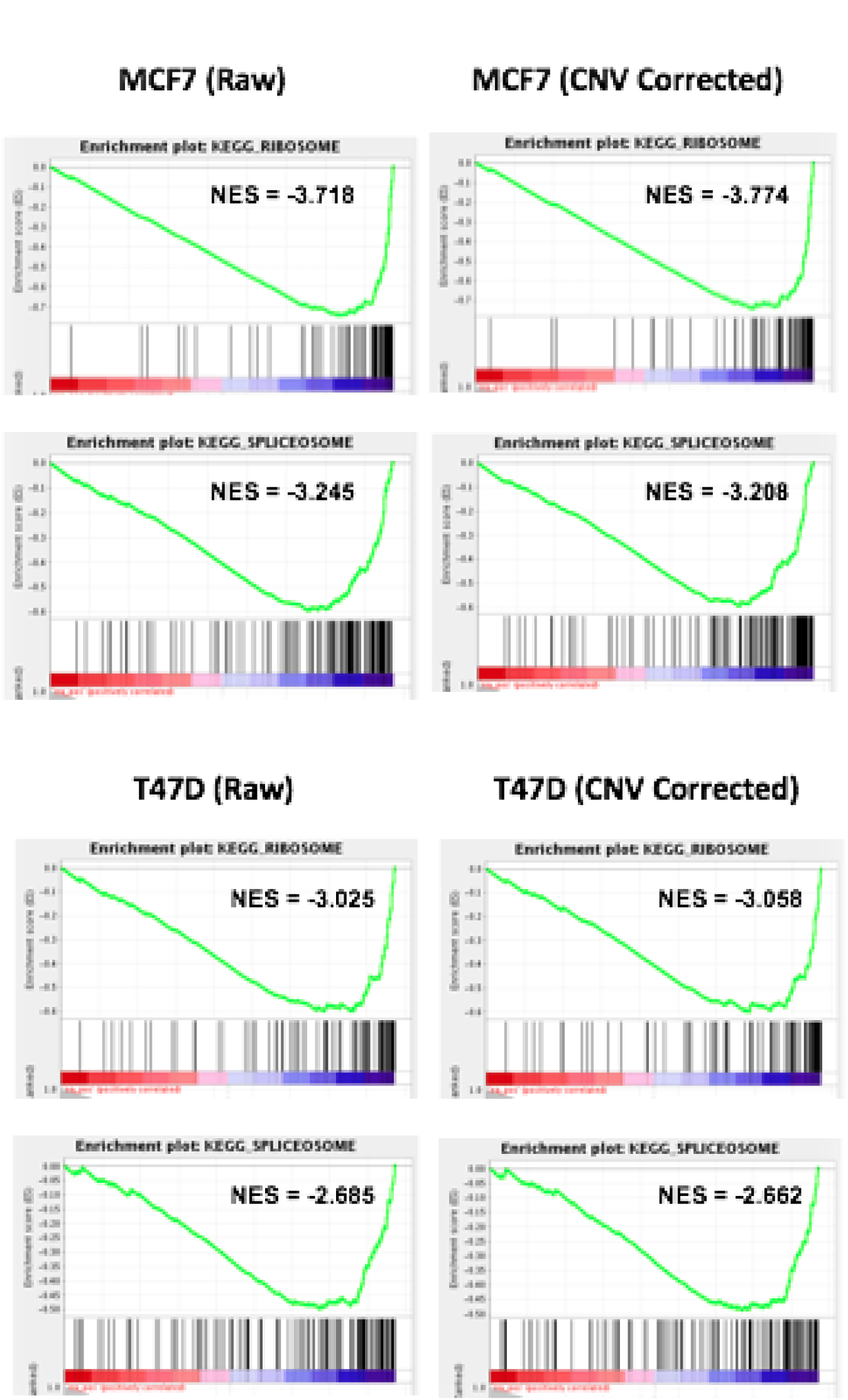
Retention of Enrichment for Highly Essential Housekeeping Genes in KEGG_RIBOSOME and KEGG_SPLIECEOSOME Pathways.

### CNV Correction without CNV profiles

For samples without CNV profiles, we employed a sliding window approach to estimate gene copy numbers by averaging the log-fold change values of sgRNAs within broad genomic windows across the entire genome. We applied this approach to MCF7 cells and compared the results with known MCF7 CNV profiles. The CNV estimates were highly correlated with actual CNV measurements in MCF7 cells, especially for regions with strong amplifications and deep deletions (Pearson’s correlation coefficient, *r* = 0.630; Figure 7a). In particular, the estimates of relative gene copy numbers in the region of Chromosome 17 correspond well to the actual CNV measurements within this same region (Figure 7b). Furthermore, when using the estimated CNV profiles for CNV bias correction, both MAGeCK-MLE and MAGeCK-RRA reduced the effects of CNV amplifications in Chromosome 17 (Figure 7c-d). Therefore, in cases where the copy number data is absent, our approach serves as a valuable tool for correcting copy number biases in CRISPR screening results.

**Figure 7.**
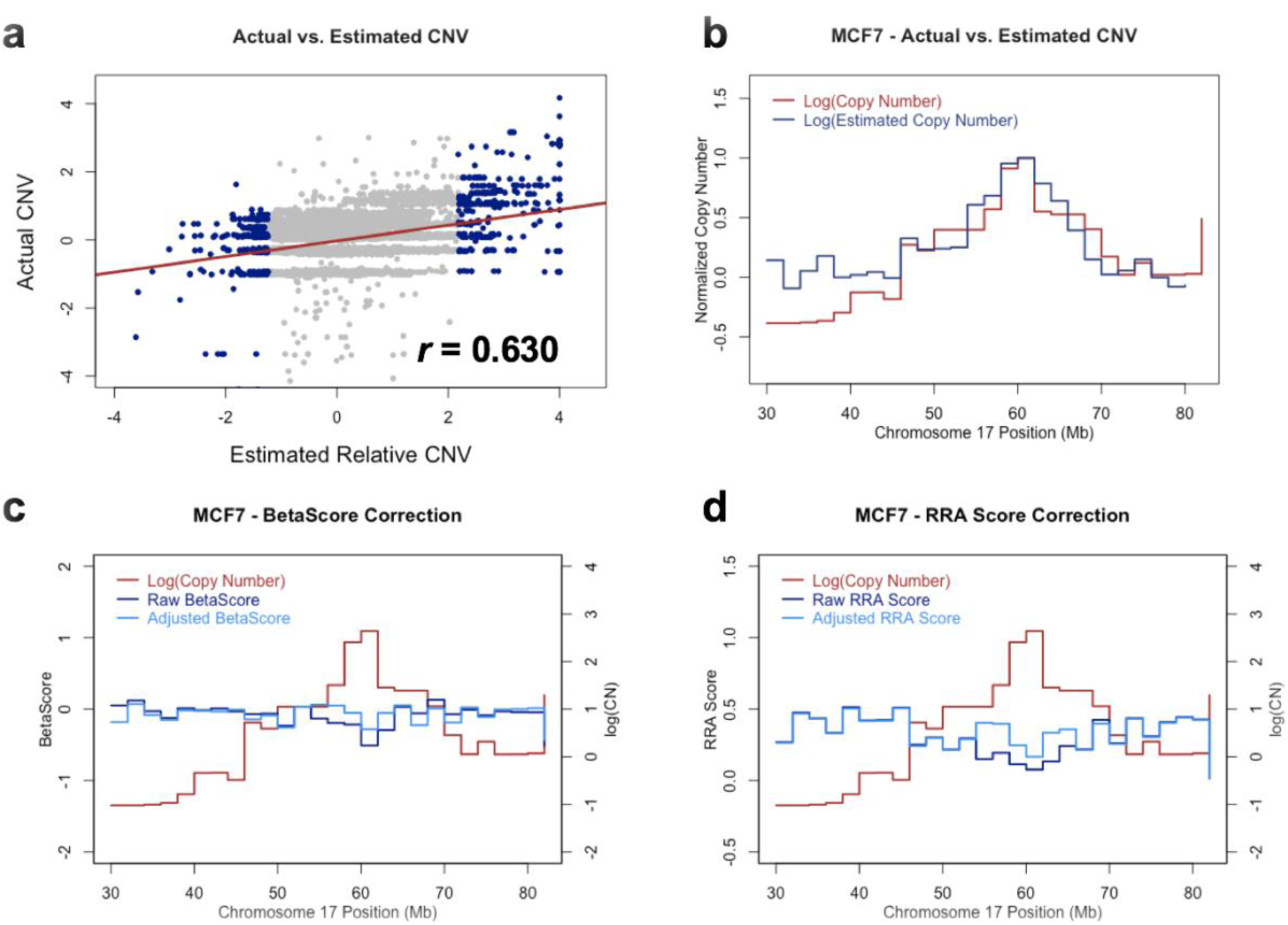
Copy Number Estimation using sgRNA Log-Fold Change. **(a)** Relationship between actual CNV values and estimated CNV values for genes with estimated CNVs in the bottom and top 2%. **(b)** Comparison of actual and estimated CNV (normalized by the max CNV value) in a high CNV region in MCF7. **(c)** *β* scores in a high CNV region in MCF7. **(d)** RRA scores in a high CNV region in MCF7.

## Conclusion

Copy number variations (CNVs) are one of the major sources of false positives in CRISPR/Cas9 knockout screens. Supplementing the current MAGeCK and MAGeCK-VISPR computational frameworks, we introduced CNV bias correction algorithms to effectively reduce biases associated with copy numbers. In addition, we developed an approach to correct CNV biases when CNV profiles are unknown, expanding the applicability of these algorithms to a broad range of CRISPR screening datasets for which CNV profiles are not available. We demonstrated that our algorithms reduced the prevalence of false positives while simultaneously retaining true essential genes, further enhancing the results of MAGeCK and MAGeCK-VISPR.

